# Comprehensive imaging of synaptic activity reveals dendritic growth rules that cluster inputs

**DOI:** 10.1101/2021.02.11.430646

**Authors:** Kaspar Podgorski, Tristan Dellazizzo Toth, Patrick Coleman, Serhiy Opushnyev, Janaina Brusco, Peter Hogg, Philip Edgcumbe, Kurt Haas

**Affiliations:** Department of Cellular and Physiological Sciences and the Brain Research Centre, University of British Columbia; Vancouver, BC, Canada V6T2B5; Janelia Research Campus, Howard Hughes Medical Institute; Ashburn, VA, USA 20147

## Abstract

The distribution of synapses across dendritic arbors determines their contribution to neural computations since nonlinear conductances amplify co-active clustered inputs. To determine whether, and how patterned synaptic topography arises during development we developed a random-access microscope capable of full-neuron calcium imaging of activity and structural plasticity of developing neurons in awake *Xenopus* tadpoles. By imaging growing brain neurons in response to plasticity-inducing visual training, we show coordinated growth and synaptogenesis specific to each neuron’s spike tuning. High evoked activity in neurons tuned to the trained stimulus induced pruning of non-driven inputs across the dendritic arbor as these neurons strengthened their responses to this stimulus. In stark contrast, initially unresponsive neurons that shifted their spike tuning toward the trained stimulus exhibited localized growth and new responsive synapses near existing active inputs. These information-driven growth rules promote clustering of synapses tuned to a developing neuron’s emerging receptive field.

**One-Sentence Summary:** Sensory input directs brain neuronal growth and connectivity promoting clustering of synaptic inputs tuned to a neuron’s encoding properties.

## Main Text

Neurons in the brain integrate inputs from hundreds to thousands of upstream neurons to perform computations, an ability that relies on the shape of their dendrites and the placement of input synapses(*1*–*3*). The spatial arrangement of dendrites determines the axons a neuron can contact, and coincident signals from nearby synapses can interact, making the placement of synapses with coincident activity important for dendritic integration(*2*–*4*). Recent studies have found evidence for spatial clustering of synapses tuned to the same input in some neuron types, and no evidence in others(*5*–*18*). It remains unknown how mature spatial arrangements of synaptic tuning arise. Patterns of coincident synaptic activity have not been measured on a large scale due to the difficulty of simultaneously measuring activity throughout complex 3D dendritic arbors in awake animals. To address this issue, we developed methods for comprehensive imaging of individual brain neurons, recording detailed dendrite growth, somatic firing, and complete patterns of 3D dendritic arbor activity driven by synaptic input. A major obstacle to comprehensive imaging *in vivo* has been the need to rapidly image large 3D volumes at high resolution. To overcome this, we developed a random-access multi-photon (RAMP) microscope(*19*–*21*) to track a neuron’s entire dendritic structure and its growth over time. RAMP microscopy enables fast high-resolution laser scanning of arbitrary sets of points in 3D volumes without taking time to image intervening space, allowing us to record the activity of all dendritic branches and the soma of a labeled neuron at volume rates ranging from 6-20Hz, depending on arbor size.

We performed comprehensive imaging of individual growing neurons in the brains of awake albino *Xenopus laevis* tadpoles while conducting plasticity-inducing visual training(*22*) to investigate how subcellular patterns of activity driven by sensory experience shape neuronal structure and function. The visual system of the transparent *Xenopus* tadpole provides *in vivo* access to a conserved vertebrate brain circuit undergoing rapid synapse formation and experience-dependent refinement(*22*–*28*). Time-lapse imaging of retinorecipient tectal neurons has revealed rich structural dynamics of dendritic filopodia(*29*), fine protrusions that act as sites of synaptic contact(*28*, *30*). Calcium indicators report tectal neuron action potential firing rates at the soma(*22*, *24*) and synaptic activity in dendrites(*8*, *23*). Visual training induces strong structural and functional plasticity in tectal neurons within 30 minutes(*22*), providing a unique opportunity to observe experience-induced plasticity in real time. We found that visual training drives patterned structural changes in dendrites, including clustered filopodial extensions and additions, and distributed retractions and subtractions. Growth and pruning responses were dependent on each neuron’s spike tuning to the training stimulus, demonstrating that experience-driven growth is dependent on alignment of the input with the encoding properties of each neuron. Using RAMP microscopy, we found that synaptic inputs are clustered along dendrites according to their tuning, and that development of this clustering is activity-dependent and predicted by local and global calcium signals. Training-induced additions were preferentially located near existing synapses activated by the training stimulus, and newly-formed synapses preferentially responded to that stimulus. Dendritic growth and pruning patterns were predicted by relative amplitudes of evoked calcium transients at the tips of filopodia and the adjacent dendritic shaft. These growth rules can explain the origin of clustered synaptic tuning and how experiences promote the encoding of specific sensory features.

## Results

### Sensory-evoked activity patterns direct dendrite growth and pruning

To correlate structural and functional plasticity in individual tectal neurons, we imaged somatic firing and dendrite growth simultaneously throughout visual training. Neurons in the intact brain were selected for labeling and imaging based on their spike tuning using targeted single-cell electroporation(*31*, *32*). After filling the tectum with the cell-permeable calcium sensor Oregon Green BAPTA 1-AM (Fig. 1a,b)(*22*), we selected a neuron based on its somatic firing response to 50ms full-field darkness (OFF) stimuli, detected by two-photon calcium imaging(*22*). The tip of a dye-filled pipette was positioned against the selected neuron’s soma and brief electrical stimulation filled the cell with Alexa Fluor 594-dextran (Fig.1b). We labeled only type 13b pyramidal neurons(*33*) in the rostral tectum. The sensory-evoked responses and extensive dendritic arbors of these neurons reflect a relatively late stage of development(*22*, *29*).

**Fig. 1.**
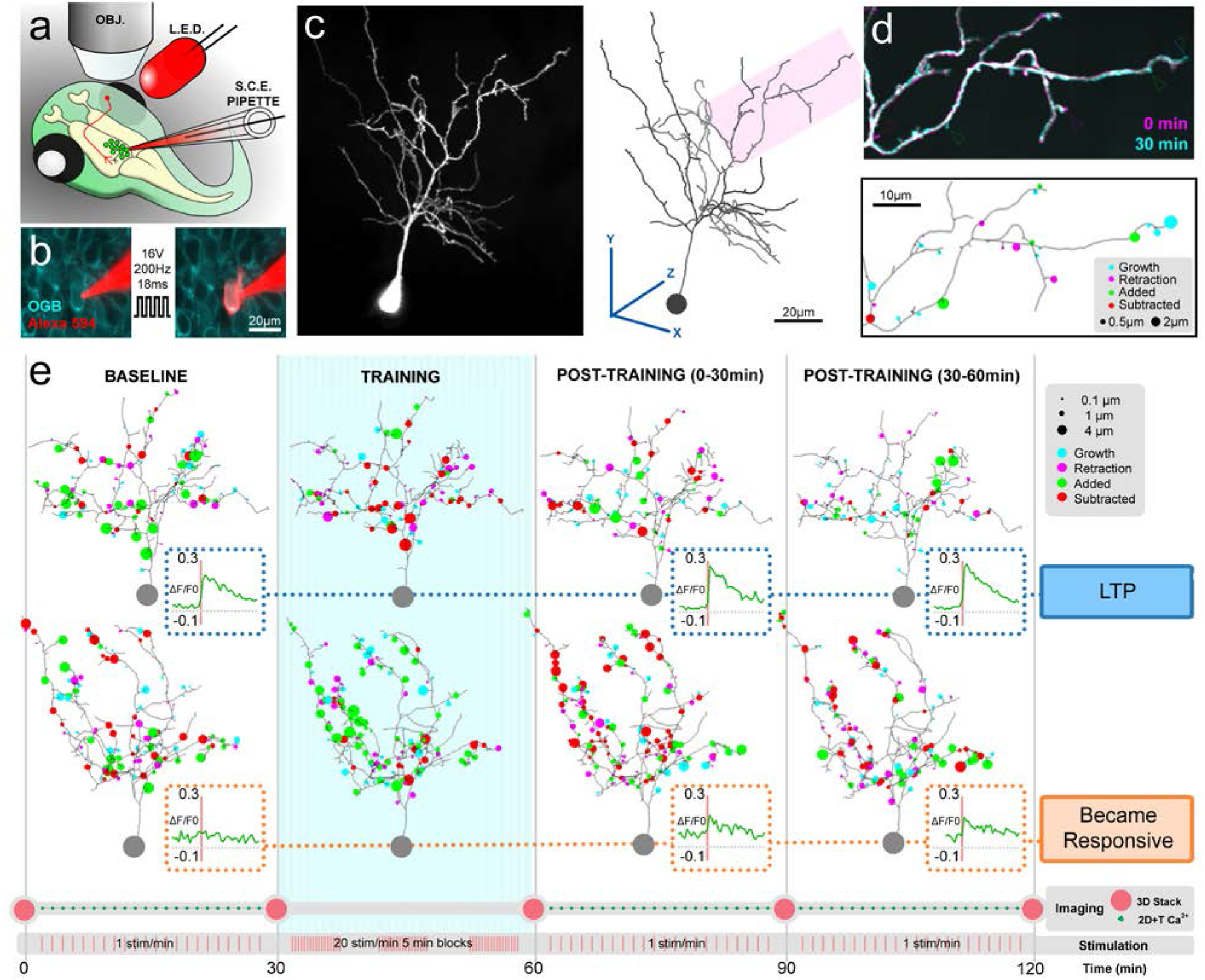
Simultaneous imaging of evoked somatic activity and structural plasticity in functionally-targeted tectal neurons. **A)** Full-field OFF visual stimuli were presented to awake tadpoles during two-photon calcium imaging of the tectum to identify responding neurons for targeted single-cell electroporation. **B)** Neurons loaded with OGB (cyan) immediately before (left) and after (right) electroporation of a single responsive neuron with space-filling Alexa 594 dextran dye (red). **C)** Z-projection (left) of a neuron image stack and corresponding 3D reconstruction (right). **D)** (top) Overlay of two substacks of the same neuron, acquired 30 minutes apart. The overlay is white where structures are stable, cyan where they extend, and magenta where they retract. Arrowheads mark sites of addition (green), subtraction (red), extension (cyan) and retraction (magenta). (bottom) Corresponding motility plot. Circles mark sites of motility, and their area represents the change in length of the filopodium. **E)** Example motility plots and mean evoked somatic responses (insets) throughout probing and training, for representative neurons showing two of the four most common activity profiles *LTP* and *Became Responsive*.

Following single-cell electroporation, we exposed tadpoles to plasticity-inducing visual stimuli to examine the effects of sensory experience on neuronal growth and pruning. We trained tadpoles using a Spaced Training (ST) protocol that primarily drives long-term potentiation (LTP) of visually-evoked somatic firing(*22*, *34*). This experiment consisted of 30 minutes of baseline probing with low-frequency OFF stimuli to assess evoked responses, followed by 30 minutes of ST with high-frequency OFF stimuli, followed by 60 minutes of post-training probing to assess plasticity(*34*). We monitored somatic activity throughout this experiment with rapid 2D imaging, and detected structural changes every 30 minutes by collecting full 3D images. Dendritic arbors were fully reconstructed in every 3D image allowing comprehensive morphological analysis of all structural changes over time (Fig. 1c,d). We grouped neurons into four activity profiles according to their initial somatic response to OFF stimuli and changes in amplitude with ST (Fig. 1e, S1a): ***LTP*** −an initial response that increased with training (n = 7), ***Stable Response*** −an initial response that did not change in amplitude (n = 5), ***Became Responsive*** −no initial response but a response after training (n = 5), ***No Response*** −no response before or after training (n = 4).

We found that each group showed distinct patterns of experience-driven structural changes (Figs. 1e, 2, S1a), with activity profiles explaining most of the variation in growth patterns (64%, MANOVA F, p<0.001). *LTP* neurons showed decreased filopodial additions during training, which remained low during post-training probing, and transiently increased subtractions during training. *Stable Response* neurons showed no significant changes in addition or subtraction rates. *Became Responsive* neurons showed a strong increase in additions during training that remained high. Subtraction rates increased strongly, but only after these neurons became responsive after training. Surprisingly, *No Response* neurons showed altered growth patterns in response to ST, with transient increased additions during training, and reduction in subtractions. These results show that increased evoked firing to the trained stimulus correlates with dendritic pruning, while lack of evoked firing correlates with growth. This is consistent with synaptic maturation in developing tadpole tectum(*28*), as well as LTP(*35*) and learning in mature rodents(*36*) associated with decreased synapse number but increased synapse size. Similarly, developing axons stabilize when their activity is correlated to postsynaptic neurons, and grow when it is not(*26*). In principle, this pattern of structural plasticity may permit neurons well-connected, and tuned to a strongly driven functional circuit to further specialize towards this circuit by eliminating non-driven inputs(*24*), while allowing neurons loosely integrated into the driven circuit to enhance these connections leading to a shift in their spike tuning.

**Fig. 2.**
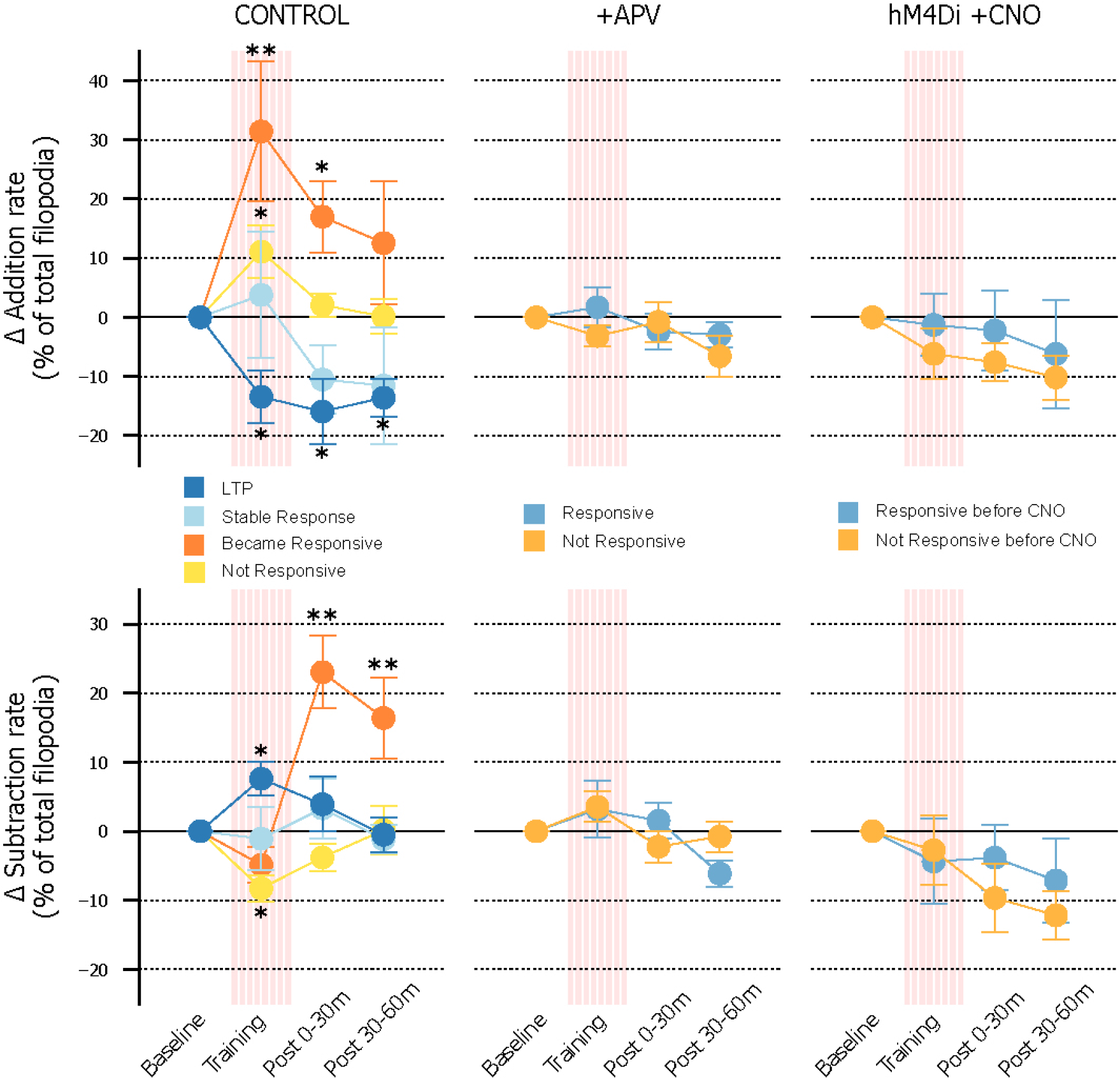
Visually-evoked activity patterns determine structural plasticity. Mean filopodial addition (top) and subtraction (bottom) rates during each 30-minute epoch, for Control neurons, neurons following D-APV injection, and neurons expressing inhibitory DREADD hM4Di following CNO injection. Control: N=7 (LTP); 5 (Stable Response); 5 (Became Responsive); 4 (No Response). APV: N=4 (Responsive); 4 (Not Responsive). hM4Di: N=4 (Responsive); 4 (Not Responsive). *:p<0.05 **p<0.01, vs. same group at baseline. ANOVA followed by Tukey LSD. Error bars indicate ± standard error of the mean (SEM).

To determine whether functional plasticity is required for training-induced structural changes, we used the NMDA receptor (NMDAR) antagonist D-APV (50μM) to block functional plasticity(*27*, *37*). Tectal D-APV injection inhibits ST-induced LTP in tectum *in vivo*(*22*) and eliminated ST-induced growth changes, regardless of somatic responsiveness (Fig. 2). Since NMDAR blockade interferes with ST-induced plasticity, but does not significantly alter neuronal firing response amplitudes, we next inhibited neural depolarization and somatic firing by expressing hM4Di, a designer receptor exclusively activated by a designer drug (DREADD)(*38*). Tectal injection of the exogenous ligand CNO (5μM) reduced evoked somatic firing of hM4Di-expressing neurons (Fig. S2). These neurons showed similar levels of filopodial additions and subtractions as non-treated neurons, but were not modulated by ST regardless of their OFF somatic responses prior to CNO administration (Fig. 2). These manipulations indicate that structural plasticity patterns depend on both somatic activity and functional plasticity.

### Dendrite growth is clustered by local intracellular signals

Dendritic integration is affected by the spatial arrangement of active synapses since nearby inputs can interact nonlinearly through voltage-sensitive conductances(*39*), and the spatial arrangement of synapses can produce emergent computations(*23*). *In vivo*, clustering of synaptic inputs based on their tuning properties or shared upstream contacts have been observed in some neurons(*9*–*11*, *13*–*15*, *18*). Furthermore, experience-driven synaptic strengthening has been found to preferentially occur in clustered inputs(*40*). This raises the question of how this clustering based on input tuning arises during early brain circuit development and whether synapse formation is spatially organized.

We performed nearest-neighbor clustering analysis of experience-driven dendritic growth dynamics, and found that ST-induced additions and extensions were more clustered than chance, while subtractions and retractions were not (Fig. 3a). To distinguish whether clustered growth is mediated by intracellular or extracellular factors, we reasoned that intracellular signals would follow intracellular distances (ID), while extracellular factors would more closely follow straight-line extracellular distances (ED) (Fig. 3b). By comparing pairs of filopodia with intracellular distance similar to their extracellular distance (ID≈ED), and those with intracellular distance much longer than their extracellular distance (ID>>ED; Fig. S3) (N = 2.5 million pairs), we found that intracellular distance predicts clustered growth (Fig. 3c), indicating that training-induced growth is driven by localized intracellular signaling.

**Fig. 3.**
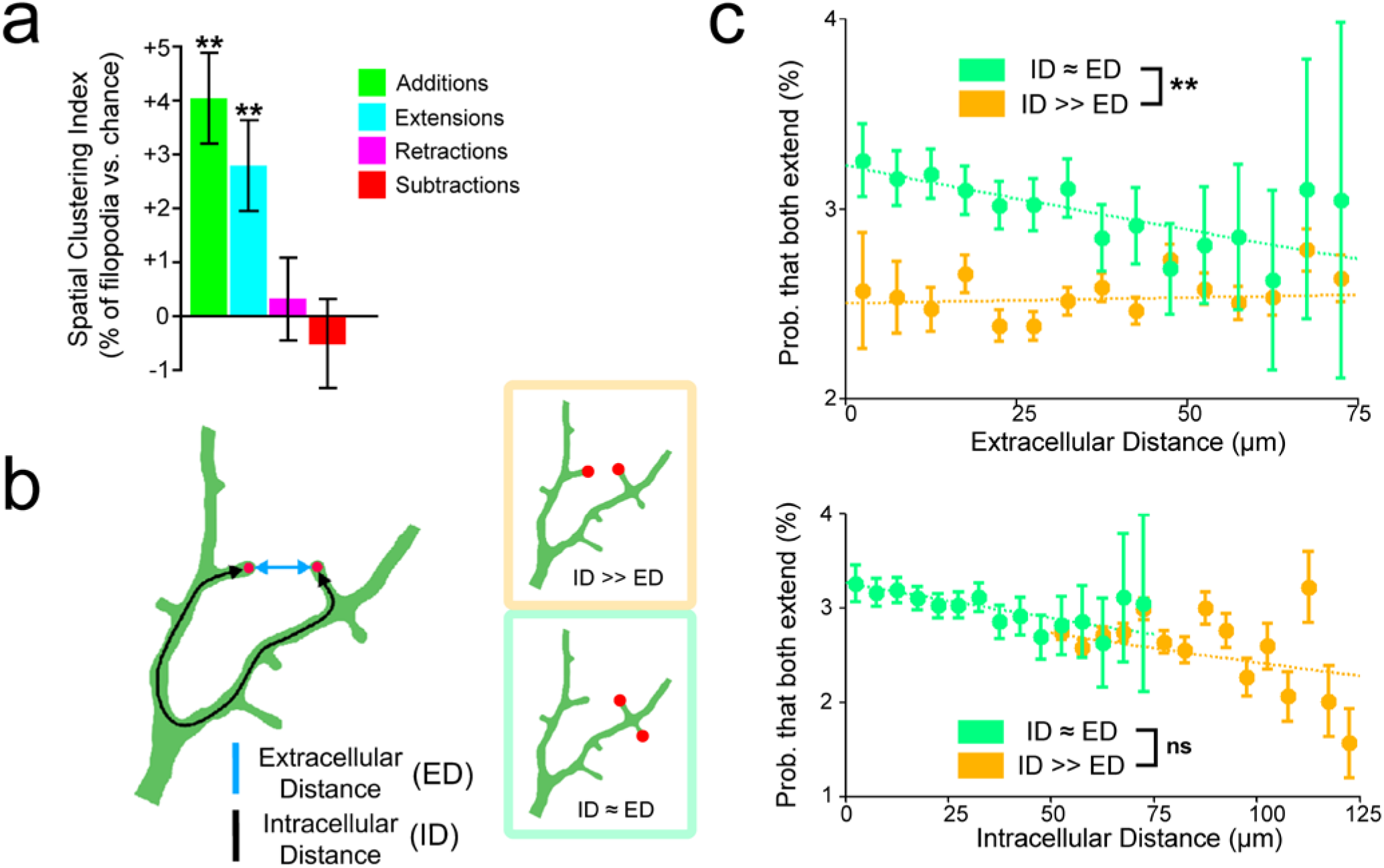
Filopodial extensions are clustered by intracellular cues. **A)** Spatial Clustering index (proportion of nearest neighbor distances less than 5 μm, minus chance proportion) for each filopodial motility type. N=1504 additions, 1102 subtractions, 1264 extensions, 1175 retractions in 21 neurons. ** p<0.01, t-test. Error bars indicate ± SEM. **B)** (left) Distances between filopodia can be measured either extracellularly (i.e. straight-line distance, blue), or intracellularly along the dendritic arbor (black). (right) We assessed whether clustering is mediated by intracellular (ID) or extracellular (ED) distance, by comparing pairs of filopodia having similar ID and ED (ID≈ED) to pairs having longer ID than ED (ID>>ED). **C)** (top) Probability that both filopodia in a pair extend, binned by intracellular distance. (bottom) Probability that both filopodia in a pair extend, binned by extracellular distance. ID predicts similarity in motility regardless of ED. ED predicts similarity in motility only when ID≈ED. Dashed lines are fit logistic regressions. ** p<0.01, Likelihood Ratio test. n=2500642 pairs in 21 neurons. Error bars indicate ± standard error of the proportion (SEP).

### Comprehensive imaging of synaptic activity, firing, and growth

The observation that growth was clustered by local intracellular cues suggested to us that spatial patterns of evoked synaptic activity could instruct the detailed shape and connectivity of a growing neuron’s dendritic arbor. However, thorough measurements of spatiotemporal input patterns across the dendritic arbor *in vivo* are not possible with conventional raster imaging techniques due to slow rates of imaging 3D volumes. To address this, we constructed a random-access multi-photon microscope(*19*–*21*) capable of imaging hundreds of dendritic sites at 6-20Hz across a 100×100×120μm volume in the intact, awake brain (Fig. 4a,b, 6a). Our microscope software performs user-assisted semiautomatic tracking of all filopodial tips, branchpoints, and all dendritic shafts (at 2μm increments) across the entire dendritic arbor of tectal neurons (Fig. 4a,b), which have 200-600 synapses predominantly localized at filopodial tips(*28*). Slight neuronal movement is accommodated by an 11-pixel sweep across each dendritic position sampled, and growth changes are accounted for by retracing the neuron’s morphology at 30-minute intervals. The tadpole model system is advantageous for RAMP imaging because brain movement from blood flow and respiration is virtually absent. By simultaneously tracking activity at all target sites as neurons grow, this method enables comprehensive imaging of both structure and function over time. We performed comprehensive imaging in single neurons expressing GCaMP6m(*8*) and jGCaMP7s(*41*). In GCaMP-expressing neurons, filopodia showed highly localized, stimulus-tuned evoked calcium transients (Fig. 4b) that were inhibited by D-APV, indicative of glutamatergic synaptic transmission (Fig. S4c,d).

**Fig. 4.**
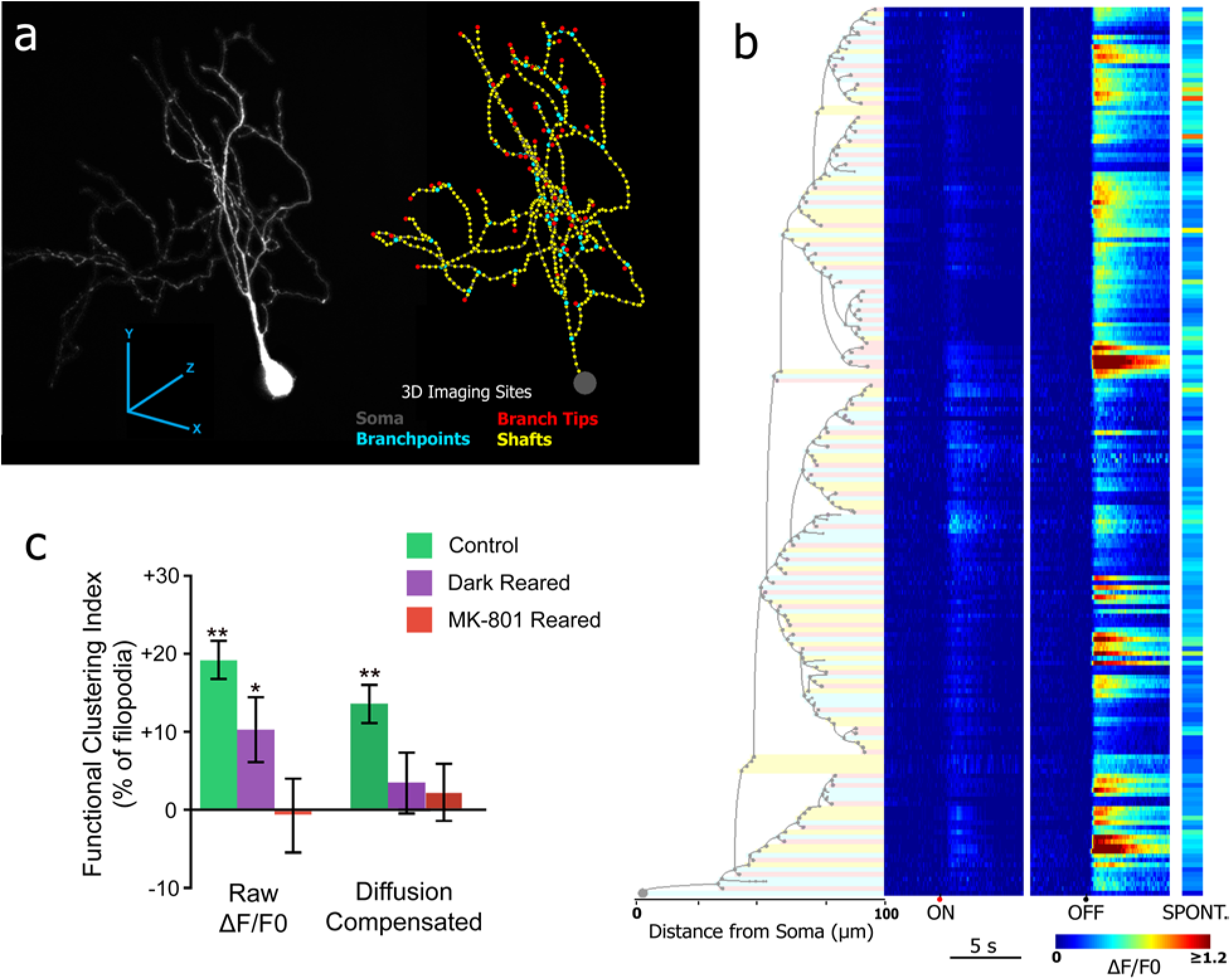
Comprehensive imaging of dendritic activity reveals spatially structured calcium signals. **A)** Maximum intensity projection (left) of a GCaMP6-labelled neuron imaged in the intact and awake brain, and corresponding wire diagram (right) with random access imaging sites overlaid. Our imaging software selects and tracks target sites at the soma, every branch tip (red), branch point (cyan), and dense samples along each shaft (yellow). **B)** Dendrogram (left) of a neuron’s structure, and ON-and OFF-evoked responses and spontaneous activity (right) at each imaging site on the neuron. Shading of dendrogram denotes the imaging site type, as in (a). Mean of n=16 responses per stimulus (ON, OFF), 3200 frames (Spontaneous). **C)** Clustering of tuned evoked tip calcium transients (left) and diffusion-compensated responses (right) in control, dark-reared, and MK-801-reared tadpoles. Diffusion-compensated inputs are clustered by their stimulus tuning in control, but not dark-or MK-801-reared tadpoles. Control: N=13 neurons. Dark Rear: N=4 neurons. MK-801: N=4 neurons. *:p<0.05, **:p<0.01, t-test. Error bars indicate ± SEM.

### Experience drives clustering of tuned synaptic inputs

To test whether synaptic tuning was spatially organized we used a modified probing protocol incorporating alternating sessions of OFF and ON stimuli, which activate different upstream retinal circuits. Evoked filopodial calcium transients showed tuning toward OFF, ON, or both stimuli. This tuning was clustered (Fig. 4b,c), indicating that sensory stimulation evokes spatially-patterned dendritic responses. This organization could be due to clustered input tuning, or to signals spreading within dendrites. To distinguish these factors, we compensated for diffusing signals at filopodium tips, reasoning that signals diffusing into the tip must pass through adjacent dendritic shaft(*8*). Diffusion compensation dramatically reduced noise correlations between nearby filopodia (Fig. S4e-g) indicating removal of signal spreading. Filopodia continued to show significant clustering of ON and OFF selectivity after diffusion compensation (Fig. 4c), suggesting that input tuning is clustered along dendrites. Supporting this interpretation, neurons of tadpoles reared in darkness or with the NMDAR blocker MK-801 (10μM) did not show clustered tuning after diffusion compensation (Fig. 4c), indicating both that diffusion-compensation effectively reduces spatial correlations in these experiments and that clustering is experience-dependent. These results supported our earlier finding that training-induced growth and pruning patterns depend on somatic depolarization and NMDAR activation, and suggest that experience-dependent growth and pruning produce clusters of similarly-active synapses.

### Local activity patterns instruct dendritic growth and pruning

We next investigated whether local patterns of sensory-evoked activity direct local patterns of structural plasticity. Local activity can induce filopodial growth or stabilization *in vitro*(*42*–*44*), as well as structural and functional plasticity in dendritic spines in mature circuits(*45*, *46*). Several types of rules are proposed to govern how calcium transients direct plasticity, based on the amplitude of calcium signals at that site(*42*, *43*, *47*), or through comparison of signals across sites, such as the correlation of a synapse to its neighbors(*9*, *48, 49*) or to somatic spiking(*27*, *50*). We evaluated the predictive power of several hypothetical rules considering both evoked and spontaneous calcium transients. Calcium transients at filopodium tips did not predict training-induced structural changes considered on their own (Fig. S5a), and correlations among filopodia or between the soma and filopodia were also not predictive (Fig. S5b,c). However, filopodia that were motile during training showed lower evoked responses at their tip compared to their base (78/92, 85%), and filopodia with larger tip than base responses were nearly all stable (350/364, 96%) suggesting that filopodia lacking strong training-evoked inputs are destabilized (Fig. 5a-d). Subtractions and retractions tended to occur in regions of higher evoked dendritic shaft transients (Fig. 5a-c). These predictive rules were specific to the training (OFF) stimulus, and did not hold for responses to the untrained stimulus (ON) or spontaneous activity (Fig. 5a), indicating that unstable filopodia do not solely represent processes lacking synapses(*30*), but that training-induced motility is sensitive to input tuning.

**Fig. 5.**
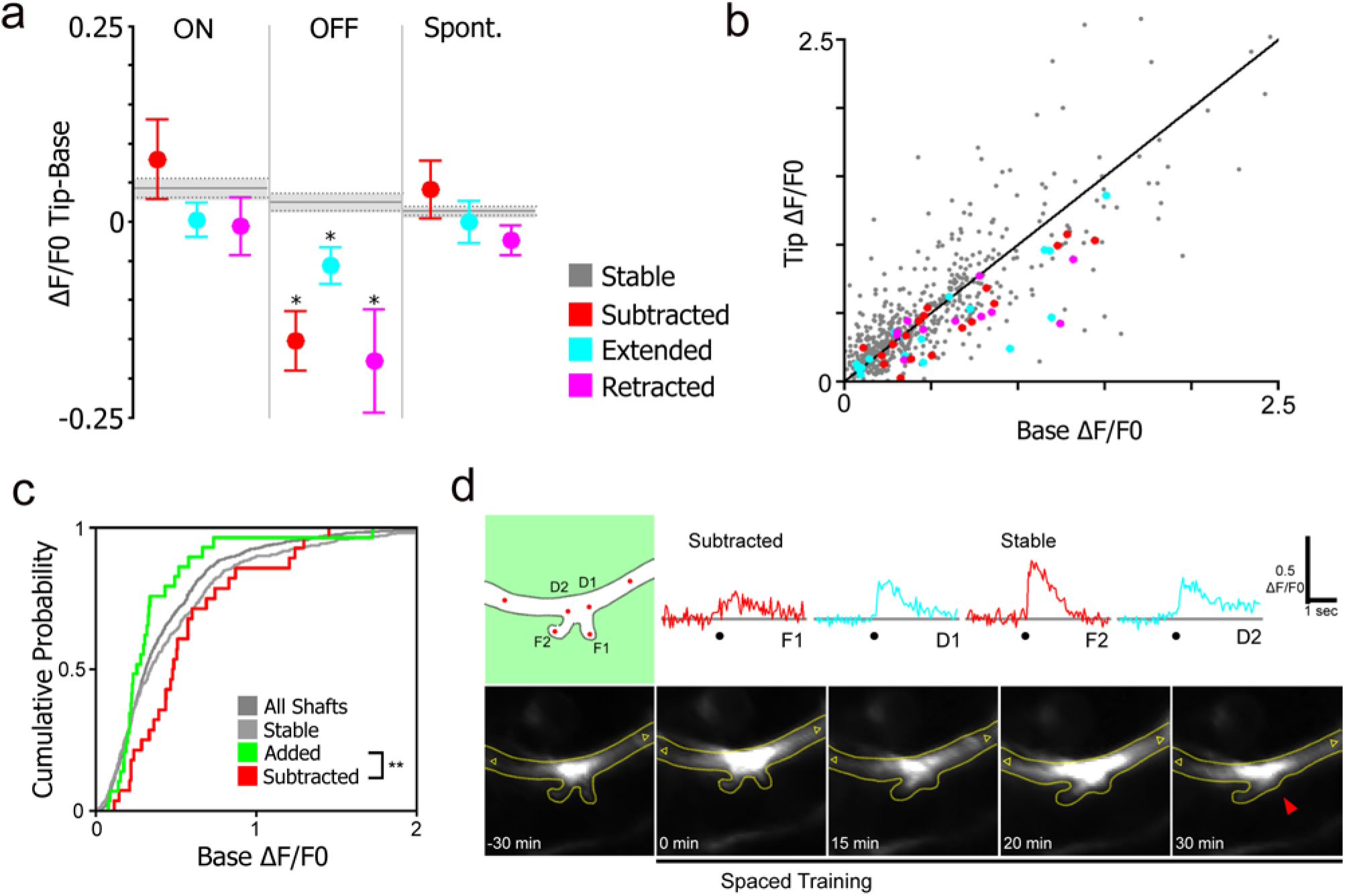
*S*patial patterns of activity predict structural changes in dendrites. **A)** Difference between tip and base calcium signals for filopodia grouped according to their growth behavior during training. Stable filopodial values are represented by the horizontal grey line in each plot. Motile filopodia showed lower tip than base transient amplitudes specific to the trained (OFF) stimulus. *: p<0.05 vs. Stable, ANOVA followed by Tukey HSD. Error bars indicate ± SEM. **B)** Mean amplitude of base and tip OFF-evoked calcium transients prior to training, colored according to that filopodium’s motility during training. N = 13 neurons, 1152 stable filopodia, 41 subtractions, 31 extensions, 25 retractions, 1972 shaft sites. (p > 0.05 for all, Tukey HSD compared to stable.) **C)** Cumulative distribution of OFF-evoked dendritic shaft transient amplitudes at sites of additions, subtractions, and stable filopodia, and all dendritic shaft sites. **:p<0.01, Kolmogorov-Smirnov test. **D)** Example correlating filopodial activity and motility. (top) OFF-evoked responses during baseline. Black dot denotes time of OFF stimulus. (bottom) Corresponding morphological images showing subtraction (red arrow) of the less responsive filopodium during training.

We next tested whether filopodial additions could be predicted by local evoked calcium activity by computing a combined spatial-functional clustering index for each filopodial addition. This value is positive for filopodia that emerge closer to existing filopodia responsive to the trained (OFF) versus untrained (ON) stimulus. Spatial-functional clustering values were larger than predicted by Monte Carlo shuffling of filopodial functional identities (Fig.6b). Moreover, new filopodia nearest to existing stable OFF-responsive filopodia showed increased survival over 1h (Fig. 6c), and new, clustered filopodia that survived for 1h were preferentially responsive to the trained stimulus (Fig. 6a,d).

**Fig. 6.**
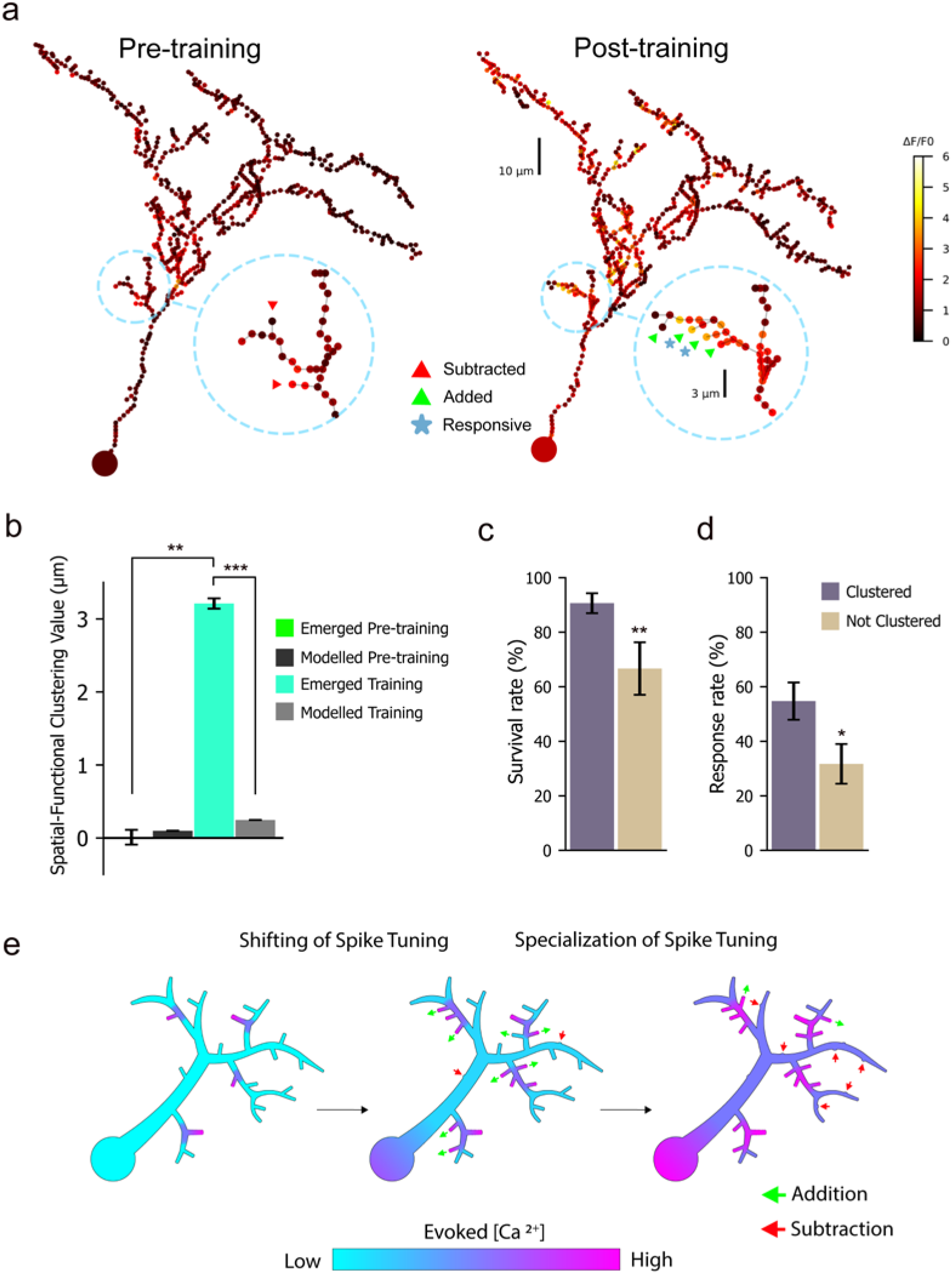
Dendrite growth behavior directed by local calcium transients promotes clustering of tuned synapses. **A)** Peak OFF-evoked responses at all imaging sites of a neuron labeled with jGCaMP7s recorded *in vivo* before (A) and after (B) Spaced Training. Inset shows filopodial additions, subtractions and responsive filopodia for the highlighted region of the neuron. **B)** Average difference in distance in microns of a filopodial addition to the nearest ON or OFF responding stable filopodia on the same neuron, termed “Spatial-temporal cluster index”. N = 14 neurons, 118 additions during training, 77 additions prior to training, 7000 Monte Carlo simulation runs to produce modelled control. *:p<0.05 t-test. Error bars indicate ± SEM. **C)** Responsiveness of filopodia added during training to the entrained stimulus. N=95 additions in 14 neurons. * p<0.05, χ^2^ test. Error bars indicate ± SEP. **D)** Survival rate of filopodia added during training over the subsequent 60 minutes. N=88 additions in 14 neurons. ** p<0.01, χ^2^ test. Error bars indicate ± SEP. **E)** Schematic of structural plasticity rules. A neuron that lacks evoked somatic firing but has sparse input from a sensory circuit (left), can respond to strong stimulus training with moderate dendritic shaft calcium near existing active synapses, which promotes clustered formation of new synapses responsive to the driven stimulus (center). Neurons that fire action potentials (spike tuned) to the driven stimulus exhibit high dendritic shaft calcium from strong local synaptic activity and back-propagating action potentials, which promote subtraction of synapses and filopodia that are not connected to the driven circuit (right). This model demonstrates how brain neurons can shift and then refine their spike tuning.

Overall, these findings show that relative levels of evoked calcium signals predict both growth and pruning patterns of dendritic filopodia (Fig. 6e). Low activity in filopodia relative to adjacent shafts predicts high mobility and subtraction, while high activity in filopodia relative to adjacent shaft predicts both their structural stability and the addition of new filopodia in their vicinity which respond to the trained stimulus.

## Discussion

We show that growth and pruning of developing dendrites is regulated by sensory experience in an activity-dependent manner based on the relationship between an environmental stimulus and each neuron’s tuning. By imaging somatic action potential activity, we demonstrate that experience-driven potentiation of somatic responses is associated with increased pruning, and our comprehensive imaging experiments show that pruning is selective for filopodia unresponsive to the training stimulus. This supports a model in which neurons well-integrated within a sensory circuit become further functionally specialized during refinement by pruning connections to less-active inputs. In contrast, neurons can shift their spike tuning to a trained stimulus by increased growth and formation of new synaptic contacts with the driven circuit until they develop somatic responses to that sensory signal. Once these neurons develop action potential responsivity to the trained stimulus, they then also specialize by pruning.

Sensory training-induced pruning was not observed in newly differentiated tectal neurons undergoing initial stages of dendritic arbor elaboration and synaptogenesis(*51*). Instead, immature tectal neurons uniformly exhibited enhanced growth in response to sensory training. During early circuit formation, sensory input without coincident firing may promote loose integration into functional circuits for the establishment of initial receptive field properties(*52*). Thus, early stimulus-driven growth mechanisms may be similar to those exhibited by more mature neurons shifting their spike tuning, and stimulus-driven pruning may emerge when neurons establish robust coordinated spike tuning.

A leading question in understanding neuronal circuit function is whether synaptic distribution across dendritic arbors is patterned, and if so, how such synaptic topography arises during neural circuit formation. Here, we find clustering of synapses with similar response properties to visual stimuli. By combining comprehensive imaging of neural activity and detailed tracking of structural plasticity we find rules based on local and global activity that drives synapse clustering based on tuning. We find that the locations of filopodial additions and subtractions are predicted by local patterns of calcium transients in the synaptic compartment and adjacent dendritic shaft. Filopodia are retracted when stimuli evoke small calcium transients in a filopodium tip and large calcium transients in the local dendritic shaft. These conditions likely reflect filopodia not receiving synaptic input when dendritic shaft calcium is high, driven by strong synaptic input and/or back-propagating action potentials(*53*). The broad distribution of backpropagating action potentials across the dendritic arbor, and the expected disparate connectivity of developing neurons to a diversity of upstream inputs are consistent with our observation that retractions are not spatially clustered. In contrast, morphological stabilization occurs in filopodia with larger calcium transients at their tip than their base. Critically, active filopodial promote the addition of new filopodia in close proximity that respond to the same stimulus, leading to spatial clustering of synapses tuned to a given stimulus.

In mature circuits, activity-driven formation of new synapses has been found to be clustered(*9*, *10*, *17*, *54*), but serial mapping of synapse tuning *in vivo* has found evidence for spatial clustering of synapses tuned to the same input in some neurons, but not others(*5*–*18*). Our results present an important distinction that could explaining these differences. Tectal neurons show clustered input tuning that is activity-dependent but stimulus-specific, due to mechanisms driving synaptic clustering only among inputs aligned with the cell’s spike tuning. Indeed, neurons using nonlinear integration to perform a computation would be expected to cluster inputs specific to that computation, and not other inputs. Such patterns may not be detectable in distributions of synaptic tuning to an arbitrary set of stimuli not aligned with an individual neuron’s spike tuning.

Our results demonstrate mechanisms directing experience-driven patterns of growth and pruning in developing brain neurons, and identify a set of rules that combine synaptic and spike activity signals to predict growth behavior throughout the dendritic arbor. These rules allow neurons in developing brain circuits to strength or shift their spike tuning in response to strong sensory input. Results demonstrate how structured organization of synaptic distribution across the dendritic arbor arises during circuit formation by mechanisms promoting clustering of synapses tuned to the emerging somatic receptive field. Since the observed patterns of neuronal growth are directed by the salience of sensory stimuli to the tuning of each synapse on a developing neurons and the neuron’s spike tuning, our results reveal mechanism of information-driven growth.

## Funding

Canadian Institutes of Health Research (CIHR) Foundation Grant FDN-148468 (KH)

## Author contributions

Conceptualization: KP, TDT, PC, KH

Methodology: KP, TDT, PC, PH, SO, JB, PE, KH

Investigation: KP, TDT, PC, PH, SO, JB

Visualization: KP, TDT, PC

Funding acquisition: KH

Project administration: KH

Supervision: KP, KH

Writing – original draft: KP, KH

Writing – review & editing: KP, TDT, PC, PH, KH

## Competing interests

Authors declare that they have no competing interests.

## Data and materials availability

All code used in the analysis is available at github.com/haaslab/dynamo. All data are available in the main text or the supplementary materials.

## Supplementary Materials

Materials and Methods

### Supplementary Text

Figs. S1 to S5

Movies S1 to S2

## Supplementary Materials

### Materials and Methods

#### Animal rearing conditions

Freely-swimming albino *Xenopus laevis* tadpoles were reared in 0.1x Steinberg’s solution (1x Steinberg’s in mM: 10 HEPES, 58NaCl, 0.67KCl, 0.34Ca(NO3)2, 0.83 MgSO4, pH 7.4) and housed at room temperature on a 12hr light/dark cycle. For MK-801 rearing experiments, tadpoles were reared in Steinberg’s solution containing 10 μM MK-801 (Tocris) from Stage 30 to Stage 50, when imaging was performed. Rearing solution was changed every 12 hours. For dark-rearing experiments, tadpoles were kept in darkness, and handled only under infrared illumination, from Stage 30 to Stage 50 (time of imaging). Experiments were conducted in accordance with the Canadian Council on Animal Care guidelines, and were approved by the Animal Care Committee of the University of British Columbia Faculty of Medicine.

#### Imaging conditions

For all experiments, tadpoles were placed in a bath containing 4mM pancuronium dibromide (a reversible paralytic) for 5 minutes immediately before imaging, then placed in a custom-fabricated chamber that immobilizes the tadpole head for awake imaging and visual stimulation. The tadpole was perfused with oxygenated 0.1x Steinberg’s solution during imaging.

#### Calcium indicator loading and tectal infusions

For population calcium imaging and targeted single-cell electroporation experiments, Oregon Green BAPTA-1 AM (OGB; Molecular Probes, Eugene, OR) was pressure injected into the left optic tectum as described previously(*24*), 30-60 min before imaging. Where APV was used, D-APV (50μM) in Ringer’s solution was injected prior to experiment onset (or included in the dye-loading solution in OGB experiments), followed by a second injection of D-APV (50μM) in Ringer’s solution prior to training onset. Control experiments in all cases consisted of the same injection protocol with drug omitted.

#### Targeted neuronal silencing

In neuronal silencing experiments, we co-electroporated GCaMP6m (from Addgene plasmid 40754, inserted into a modified pN1 vector, 3μg/μl) at Stage 35-36 and the DREADD HM4Di (3μg/μl) by ventricular electroporation(*55*) at Stage 35-36. Control tadpoles were electroporated with GCaMP6m (3μg/μl) alone. Tadpoles were screened for expression of GCaMP6m in isolated neurons at stage 50. For these experiments, we acquired an additional probing epoch of activity data (‘Baseline’) prior to our stimulation protocol, to assess response amplitudes prior to CNO administration. Following the Baseline epoch, tadpoles were anaesthetized and 5μM CNO was infused into the tectum. The normal stimulation protocol (see below) was initiated following recovery from anesthesia. During this imaging, the imaging chamber was perfused with Steinberg’s solution containing 0.5μM CNO.

#### Target neurons

For all experiments, we targeted type 13b pyramidal neurons(*33*) in the dorsolateral tectum of Stage 50 tadpoles. In this target region at this developmental stage, these neurons show robust visually evoked responses and are relatively stable morphologically, with motility rates that can be accurately captured by imaging at 30-minute intervals. Nevertheless, these neurons also robustly show both structural and functional plasticity with training.

#### Conventional two-photon imaging

Experiments not involving random-access imaging were performed with a galvo-based two-photon microscope adapted from an Olympus FV300 confocal microscope (Olympus) and a Chameleon XR Ti: Sapphire laser (Coherent). Population calcium imaging (Fig. S1b,c) was performed at 5Hz, as described previously(*24*). Single cells and their neighbors within a 28×28μm field of view were imaged at 6.6Hz (Figs. 1b,e, S1a).

#### Visual stimulation protocol

Visual stimuli were presented to the eye contralateral to the imaged tectum via either an LED or a projector. Where OFF stimuli were presented, the image source, (on at start of trial) was turned off for 50ms. Where ON stimuli were presented, the image source (off at start of trial) was turned on for 50ms. In all cases, Spaced Training (ST) consisted of three 5-minute bursts of high-frequency (0.3Hz, 50ms) OFF stimuli spaced by 5-minute periods of invariant light. For conventional two-photon imaging experiments, visual stimulation consisted of three probing epochs of 30 minutes each (Fig. 1e, S1a), during which we presented one OFF stimulus per minute while recording somatic activity of the target neuron and its neighbors. Probing was halted during morphological imaging, for a 5-minute period between each epoch. ST was performed between the first and second probing epoch, for a total stimulation period of 2 hours. Similarly, visual stimulation for random-access imaging consisted of three 30-minute probing epochs, with OFF ST presented following baseline, for a total stimulation period of 2 hours. Each probing epoch consisted of at least 8 probing sessions. During each probing session, 4 stimuli were shown, consisting of either ON or OFF flashes presented with pseudo-random inter-stimulus intervals ranging from 8-12 seconds. ON and OFF probing trials were alternated throughout the experiment. During full 3D neuronal morphological imaging, registration, and acquisition planning (~2 minutes between each probing session; see software section, below), the visual stimulus was turned on and off at pseudo-random intervals of 5 to 10 seconds, to ensure roughly equal amounts of light and darkness over time.

#### Processing of calcium imaging data

For conventional calcium imaging experiments, image registration, ROI selection, spatial filtering, and baseline fitting were performed as described previously to obtain ΔF/F_0_ fluorescence traces(*24*). Initial data processing for random-access imaging is described separately below. To measure response amplitudes at each dendritic or somatic site, ΔF/F_0_ traces were filtered using non-negative deconvolution(*56*) (NND) and the mean of the unfiltered trace over 0.5 seconds prior to the stimulus was subtracted from the peak of the NND filtered trace within 2 seconds after the stimulus. A decay time constant of 0.75s was used for filtering. This constant is shorter than the ‘off’ time constant of GCamP6m and was selected to avoid oversmoothing, while reducing high-frequency noise. P-values for each response were calculated as the probability of observing a peak value of the measured size in a filtered trace of Gaussian noise with the same variance as the ΔF/F_0_ trace over 2.5 seconds prior to the stimulus. Mean responses were calculated as the mean of the unfiltered ΔF/F_0_ trace over all stimuli of a given type (ON or OFF) in an epoch. Neurons were classified as ‘responding’ in a given epoch if the peak of their mean response differed significantly (p<0.05) from noise, P-values calculates as above.

For all experiments, neurons responsive during baseline were classified as undergoing ***LTP*** if their response amplitude significantly increased (p<0.05, t-test); ***LTD***, if their response amplitude decreased (p<0.05); or ***Stable Response***, if response amplitudes did not change significantly, over both 0 – 30 min and 30 – 60 min following the end of ST. Neurons not initially responsive during baseline probing were characterized as ***Became Responsive***, if they were responsive in the 0-30 min period post-ST, or ***No Response***, if they were not responsive for any epoch. Neurons not falling into any of these categories were labelled **Other**. Where activity or plasticity were blocked with hM4Di+CNO or APV, respectively, neurons were classified according to their OFF responses measured prior to baseline, as ***Responsive*** (peak ΔF/F_0_>0, p<0.05, t-test) or ***Unresponsive***.

#### Targeted Single-Cell Electroporation

For dual somatic activity and full 3D morphology sampling, neurons were labelled by targeted single-cell electroporation (TSCE) using a novel targeted approach based on a neuron’s spike tuning after bolus loading of OGB. Using two-photon imaging of OGB fluorescence responses evoked by the visual stimulus, we select a neuron on the basis of its response and approach it with a pipette containing Alexa Fluor 594-dextran, which is driven into the cell using a brief train of voltage pulses (Fig. 1b). Typical electroporation parameters for dye-loading were 18ms train duration, 16V, 200 us/pulse, and 200pulses/s. We targeted either responding neurons or non-responding neurons adjacent to responding neurons in the dorsolateral tectum. Electroporation caused an immediate increase in somatic OGB fluorescence and transient loss of neuronal responsiveness that returned to pre-electroporation levels within 20 minutes. To determine the effects of electroporation on neurons, we compared stimulus-driven growth, activity and plasticity in neurons immediately after TSCE labelling, neurons labelled one day prior, and non-electroporated neurons (activity imaging only). We measured ST-induced functional plasticity in both the electroporated target neuron and its nearest neighbors loaded with OGB. Electroporated and neighboring non-electroporated cells showed no differences in plasticity outcomes with ST (Fig. S1b,c). We compared comprehensive morphometric measures in neurons labelled 30 minutes before the experiment and neurons labelled one day earlier by ‘shadow TSCE’(*31*). Neurons electroporated 30 minutes before the experiment, neurons electroporated one day earlier, and neurons labelled by ventricular electroporation of GCaMP6m at Stage 35-36, and subsequently screened to find isolated expressing type 13b neurons, showed indistinguishable growth patterns of total dendritic branch length (TDBL), filopodial additions, and subtractions (Fig. S1d). Altogether, tectal neurons appear to exhibit normal evoked responses and undergo normal experience-induced plasticity and structural growth after electroporation.

#### Single-Cell Electroporation of Genetically Encoded Calcium Indicators

To image synaptic and action potential (AP) activity in individual tectal neurons we employed single-cell electroporation(*32*) to express the genetically-encoded calcium indicator (GECI), jGCaMP7s (*41*)(*20*). A membrane localized version of GCaMP6m or jGCaMP7s was co-expressed with a membrane-localized version of the red fluorophore mCyRFP1(*57*) using a plasmid containing a self-cleaving P2A(*58*). mCyRFP1 was used as a space-filler for tracking dendritic morphology. Electroporation parameters were 1.1s train duration, −40V, 1ms/pulse, and 200pulses/s. Neurons were screened for expression after 48 hours and imaged 72 hours post-electroporation. In some experiments, neurons were labeled using whole-brain electroporation(*55*).

#### Morphological imaging and dynamic morphometrics

All morphological imaging, for both conventional and random-access microscopes, was performed by 3D raster scanning with a Z-axis step of 1.5 μm. 3D raster scans performed by the random-access microscope were completed in <1 min. Neuron tracing and dynamic morphometrics analytics were performed using our software Dynamo (github.com/haaslab/dynamo). Image stacks were denoised with CANDLE(*59*) using the following settings: smoothing parameter, 0.8; patch radius, 1; search radius, 3. Filopodia were defined as any unbranched process shorter than 10μm(*29*). Processes longer than 10μm were classified as branches. The axon and axonal processes were excluded from analysis. Only interstitial filopodia were included in clustering analyses, and were defined as filopodia originating more than 5μm from a branch tip in order to exclude those associated with dendritic growth cones. Filopodia were characterized as extending or retracting if their measured length changed by 0.5μm or more across 30 min epochs, and stable otherwise. All analyses were performed using Matlab and Python.

#### Spatial clustering

To measure spatial clustering of growth behavior and evoked responsivity, we computed the nearest neighbor distance (NND) for each property, *i.e*. the distance to the nearest filopodium with the same type of motility or tuning. For example, the NND for a filopodium added at a given time-point is the distance to the base of the nearest other filopodium added at that time-point. We excluded filopodia at the tips of branches from clustering analyses due to their association with growth cones and distinct growth patterns(*29*). To measure clustering, we compared both the proportion of NNDs less than 5μm and the average NND distance to the proportion and average expected under the null (non-clustered) distribution, obtained by Monte Carlo sampling. For properties of existing filopodia (such as subtractions, growth, retraction, or response amplitude) the null distribution was calculated by randomly reassigning observed values among all existing filopodia and measuring the NND on this randomized set. The null distribution for additions was calculated differently, by uniform randomly reassigning the observed number of addition sites along the length of the dendritic arbor. These reassignments were performed 10000 times for each neuron to generate the null distribution for each quantity. The term ‘spatial clustering index’ represents the observed proportion of NNDs less than 5μm minus the expected proportion. Error bars denote standard error of the proportion, from observed data and Monte Carlo sampling, propagated through the above calculations.

#### 2D synaptic imaging

2D imaging of localized dendritic calcium transients was performed on a conventional raster-scanning two-photon microscope. We imaged dendritic segments of mature tectal neurons in stage 50 tadpoles at 930nm excitation. Short segments with filopodia visible in the focal plane were imaged at 10Hz while visual stimulation switched between on and off at a mean interval of 8s. After this initial imaging period, tadpoles were removed from the imaging chamber and the tectum injected with 50μM D-APV in frog Ringer’s solution, or Ringer’s solution alone, and a second round of imaging was performed. Movies and corresponding individual frames of 2D synaptic imaging were produced by fitting ΔF/F_0_ for each pixel in the frame, after rigid body image registration.

#### Random access microscope design

To image localized dendritic activity, we constructed a random-access microscope. Lateral (X-Y) scanning was performed with a pair of crossed large-aperture (9mm) acousto-optic deflectors (AODs; Isomet OAD1121-XY). Prior to scanning, the excitation beam was collimated and expanded through coupled achromatic doublets, and its polarization rotated to optimize diffraction by the AODs. The diameter of the beam entering the AODs was controlled by an aperture (A1). Combined with interchangeable achromatic doublets acting as zoom lenses, adjustment of this aperture allows trade-offs between AOD access time, scan range, and focal spot size. In dendritic imaging experiments A1 was set to 6mm, producing an 8mm spot fully filling the back aperture of the objective (LUMFLN 60XW, 1.1NA, Olympus), a lateral scan range of 110×110μm, and an access time (time taken for the acoustic wave in the AOD to fully cross the excitation beam) of approximately 10μs. We used a Ti:Sapphire excitation laser with integrated tunable dispersion compensation (Chameleon Vision II, Coherent) to compensate for temporal dispersion of ultrafast pulses introduced by the AODs. All random-access imaging of GCaMP expressing neurons was performed at 910 nm excitation. Angular dispersion introduced by the AODs was compensated using a prism in the optical path immediately after the deflectors(*60*) oriented at 45 degrees to the two deflection axes. The apex angle of the prism was selected to best compensate for angular dispersion in the center of the field of view. A small portion of the excitation beam is reflected by a coverslip to a photodiode (UDT Sensors, PIN-10D) prior to entering the objective, allowing us to calibrate the intensity of the diffracted beam across the field of view for spatially uniform excitation, and to compensate for laser intensity fluctuations during imaging. Axial scanning of the excitation beam was produced by a piezo objective mount (PiezoJena, MIPOS 100PL). Emitted fluorescence was detected with H7422-40 PMT modules (Hamamatsu Photonics, Japan). PMT signals were amplified with SR570 transimpedance amplifiers (Stanford Research Systems, Sunnyvale, CA). Hardware was controlled by, and signals were acquired on, two PCI-6110 DAQ boards (National Instruments, Austin, TX), at a rate of 5MHz. PMT signals were deconvolved with non-negative deconvolution to recover pixel intensities(*56*).

#### Microscope software

All random-access imaging was performed using openly available imaging suites in Matlab and LabVIEW. We recorded comprehensive dendritic activity from single neurons in the tectum labeled with GCaMP. After collecting an initial image stack, Dynamo was used to trace the neuron. Imaging sites are automatically selected at every dendritic branch and filopodium tip, every branch point, and typically at 2 μm intervals along the entire dendritic arbor. Activity at these sites was recorded simultaneously at 6-20Hz in 40-second probing sessions (see Visual Stimulation, above). We obtained a full morphological image stack between each session, to compensate for drift or growth, and track morphological plasticity (Fig. S4b). For each 40-second probing session, our microscope controller optimizes the range and profile of the objective Z-axis scan to match the time spent in any given imaging plane to the number of points to be imaged in that plane. Each target point is sampled 5 times during each oscillation, including both its rising and falling phase. During fast scanning, each site was imaged as a short 11-pixel line scan 2.7 μm in length, allowing detection of drift and sample motion. Recordings that showed drift or motion artifacts were discarded from the time of motion onwards. In our paralyzed tadpole preparation, only a small fraction (<10%) of recordings showed detectable drift over a single imaging cycle, which lasts roughly 2 minutes.

Morphological image stacks obtained between probing sessions were used for automated registration. Registration consisted of initial global registration followed by registration of local regions (~5μm in each direction) surrounding each branch tip and branch point. Intermediate points were inferred using registered branch points as landmarks. All registrations performed were rigid-body translations based on image stack cross-correlations to the previous morphological image stack, typically acquired roughly 2 minutes earlier. Where the error between registered images exceeded a threshold, corresponding points were selected manually. Newly added filopodia were manually traced at the start of each new epoch (every 30 minutes) for inclusion in comprehensive imaging.

#### Processing of random-access imaging data

Following recording, correct registration of imaging sites to the neuron’s structure in each session was confirmed by manual examination. Each line scan was registered across frames of the fast scan to detect drift or sample motion, and each movie was further manually inspected for drift. Traces that showed drift of more than 2 pixels were discarded from the time of drift onwards. Drift was rare, with less than 10% of all traces showing more than 2 pixels of drift over 40 seconds. Raw fluorescence traces for each imaging site were generated by summing intensity of the center 7 pixels of each aligned 11-pixel line scan. ΔF/F_0_, response amplitudes, and plasticity metrics were obtained from these raw fluorescence traces in the same manner as for conventional calcium imaging data.

#### Compensation of spreading calcium signals

To compensate for calcium signals spreading into filopodium tips from the dendritic shaft, so as to isolate signals originating within filopodia, we adapted a technique used to compensate for back-propagating action potentials in mouse cortical neuron spines(*11*). The joint distribution of base and tip ΔF/F_0_ amplitudes typically shows two arms (Fig. S4f) or a fan shape, but always has a clear lower limit on the ratio of base to tip fluorescence. We attribute this lower limit to some proportion of calcium signals at the base that enter the tip by diffusion or similar processes. We fit this lower limit (the mixing coefficient) for each filopodium, for each probing session, using robust regression (function robustfit in the Matlab statistics toolbox) for data frames where base ΔF/F_0_ exceeded tip ΔF/F_0_. Compensated tip signals were calculated as (tip ΔF/F_0_) – (mixing coefficient)*(base ΔF/F_0_) and NND filtered with a 0.75s time constant. Diffusion compensation dramatically reduced locally-structured noise correlations (Fig. S4g), indicating that spreading signals were effectively removed.

#### Spontaneous activity

“Spontaneous activity” refers to fluorescence traces recorded during periods between stimulus presentations. All activity recorded from 5 seconds following each stimulus presentation up to the frame before the following stimulus was included in analysis of spontaneous activity. Mean spontaneous activity (Fig. 4b) is the mean ΔF/F_0_ value for a given site over this time.

#### Measurement of correlations

Correlations were calculated as the Pearson correlation between traces for each pair of sites on the neuron. To obtain noise correlations, correlations in evoked activity were calculated separately over OFF and ON trails then averaged, thus removing the effect of stimulus identity on the correlation. Prior to calculation of noise correlations, global variations in evoked responses (*e.g*., due to run-down of responses to repeated stimulation) were removed by subtracting the reconstructed signal produced from the largest singular value of the response matrix. For spatial analysis of diffusion compensated noise correlations, correlations between filopodia sharing a measurement point at their base (i.e. filopodia with a distance of 0) were not included in analysis, to avoid including spurious correlations due to shared variation in the base data used for compensation. Correlations in calcium signals and diffusion-compensated inputs (Fig. S5b,c) are total correlations, not noise correlations; they include variations due to the identity of the stimulus, and reflect the overall similarity in calcium fluctuations at the two sites being correlated. In these figures, ‘Correlation to nearby filopodia’ refers to the mean correlation to all filopodia pairs at intracellular distances less than 10μm.

#### Clustering of local dendritic responses

We used diffusion compensated signals to identify the tuning preferences of individual filopodia (see Diffusion Compensation). To assess clustering of ON/OFF tuning of filopodial responses, we calculated the functional clustering index by measuring NNDs separately for each stimulus, over filopodia with mean diffusion-compensated responses to that stimulus exceeding a threshold. The threshold for selection was the 80^th^ percentile of evoked response amplitudes for each neuron, bounded above and below at 0.2 and 0.6 ΔF/F_0_.

#### Image processing

For large-scale maximum intensity projections (Fig. 1c,d, 4a), regions more than 2 microns away from neuronal processes (as determined by Dynamo) were dimmed by a factor of 10, to make it possible to see small processes throughout the entire depth of the image.

#### Statistics

Statistical tests are reported alongside p-values. Matlab and Python code for Monte Carlo simulations and statistical tests are available by contacting the authors.

**Fig. S1.**
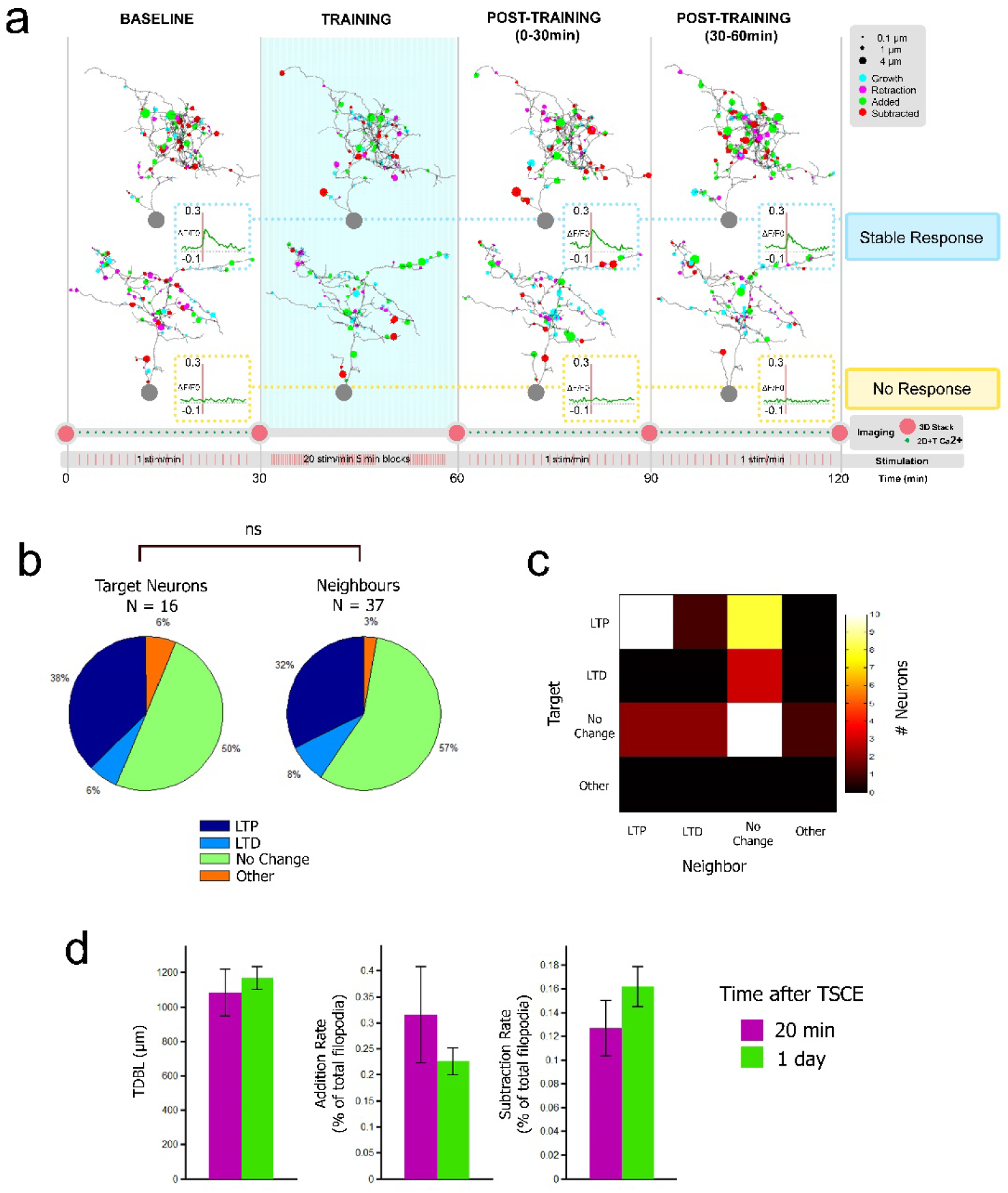
**a)** Simultaneous monitoring of evoked somatic activity and structural plasticity in neurons labeled according to visually-evoked responses. Dynamo motility plots and mean evoked somatic responses (insets) throughout probing and training, for representative neurons showing Stable Response and No Response. **b-d)** Neurons electroporated by targeted single-cell electroporation (TSCE) show normal patterns of activity and growth. (b) Distribution of plasticity outcomes to OFF spaced training for OFF-responsive neurons electroporated with TSCE and OFF-responsive unelectroporated neighbors in OGB-filled tecta. Electroporated neurons show the same distribution of plasticity outcomes as neighbors. (p > 0.05, Pearson’s χ^2^, N shown). (c) Plasticity of neighbor cells compared to plasticity of the target cell. (d) Total dendritic branch length (TDBL), filopodial addition, and subtraction rates, measured 20 minutes, or 1 day after TSCE. Neurons show normal patterns of growth measured soon after labelling. (p > 0.05, t-test, N=22 neurons). Error bars indicate ± SEM.

**Fig. S2.**
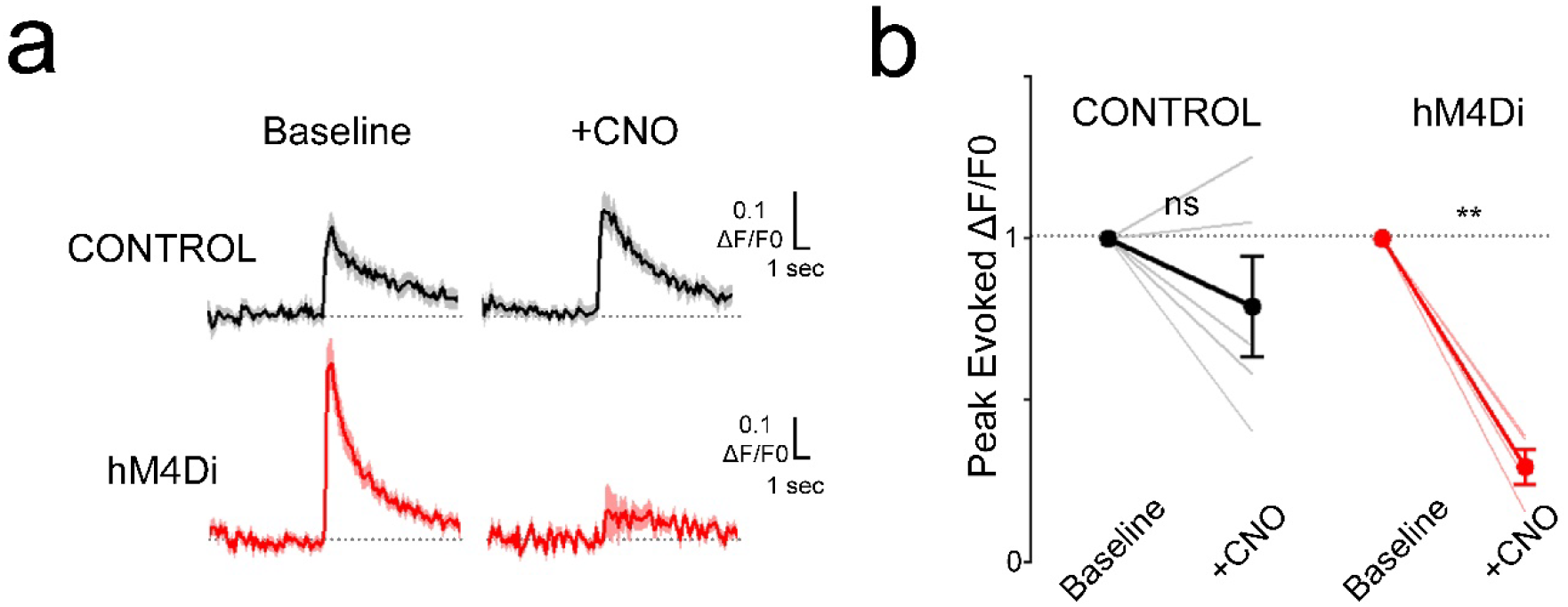
**a)** Mean responses of a neuron expressing GCaMP6m alone (Control, top) or GCaMP6m+hM4Di (bottom) before and after CNO administration. Shading denotes SEM (n=16 trials/trace). **b)** Average response amplitudes before and after CNO administration of all initially-responding Control and hM4Di-expressing neurons tested. (Control, N=5; hM4Di, N=4 neurons). **:p<0.01, paired t-test. Error bars indicate ± SEM.

**Fig. S3.**
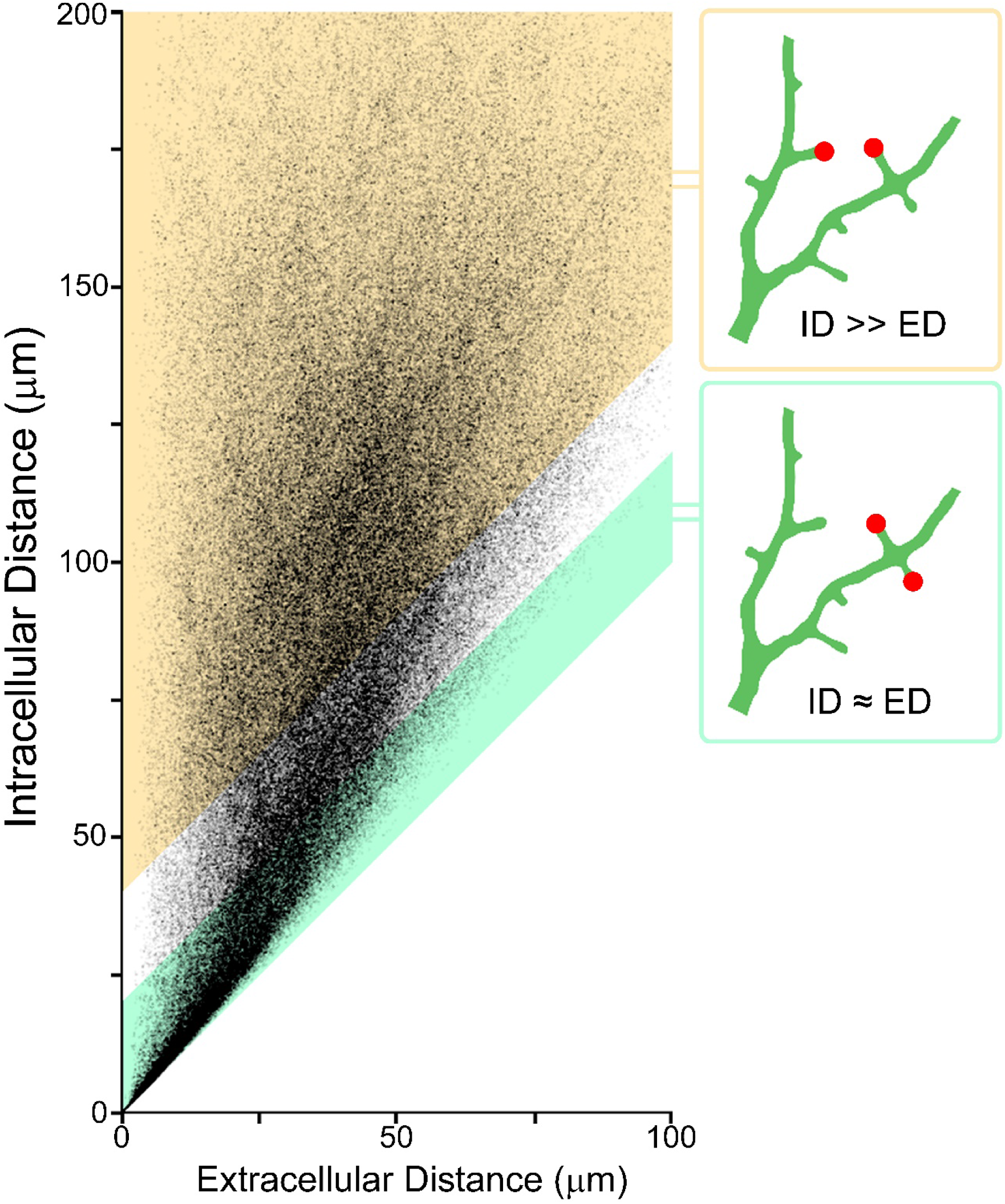
We assessed whether clustering is mediated by intracellular (ID) or extracellular (ED) distance, by comparing pairs of filopodia having similar ID and ED (ID≈ED) to pairs having longer ID than ED (ID>>ED).

**Fig. S4.**
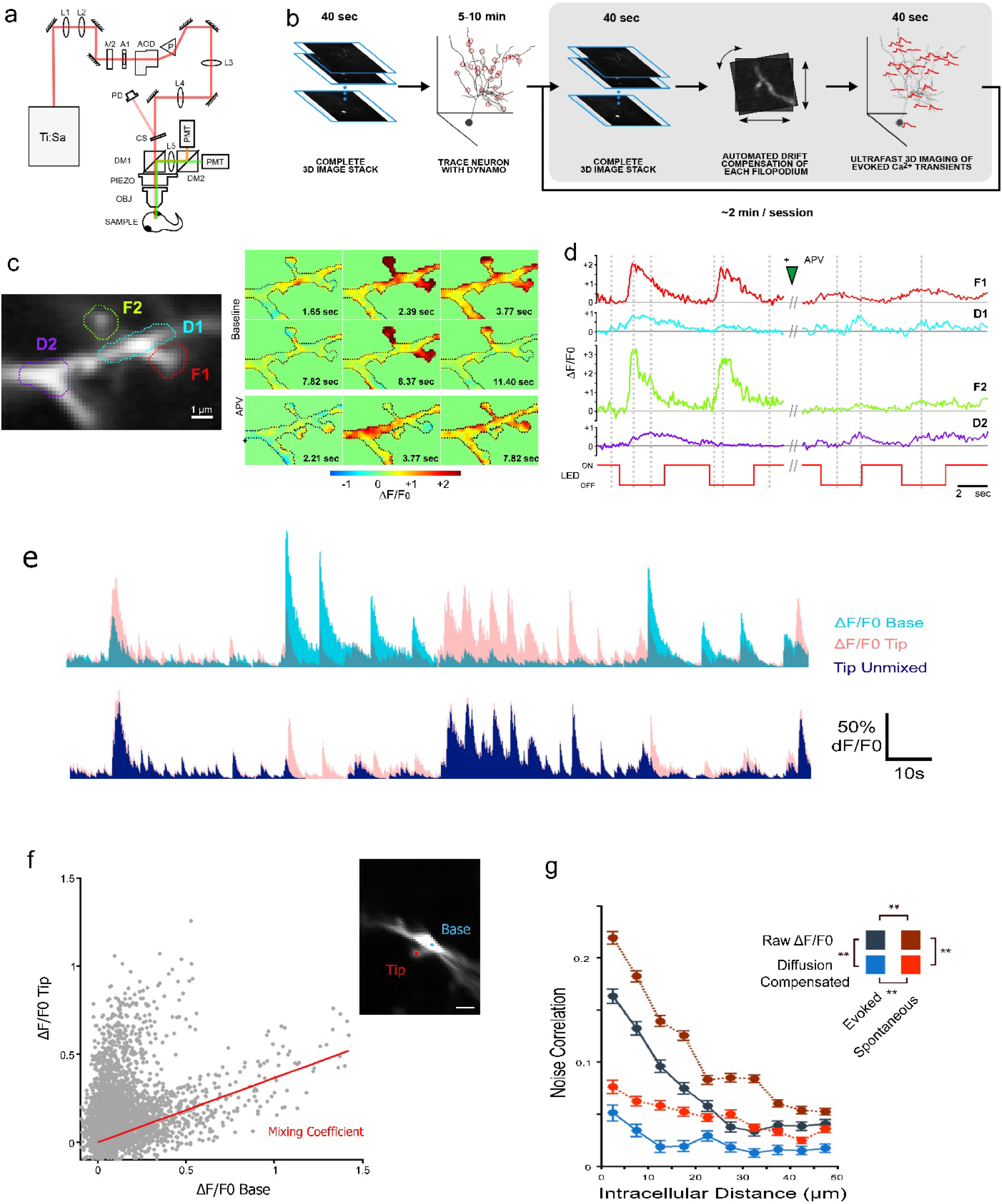
**a)** Schematic of our custom-built random-access two-photon microscope. **b)** Schematic of random-access imaging protocol. An initial volume image was acquired and traced with Dynamo for identification of imaging sites. Random access imaging consisted of alternating morphological imaging to compensate for movement and growth, and 40-second probing sessions to record comprehensive dendritic responses. **c)** Filopodia show localized, NMDA-receptor dependent sensory-evoked calcium transients. Raster-scanned image of a dendritic segment of a GCaMP6m-expressing neuron. Regions of interest for traces in are outlined. Average of 140 frames. (*right*) Individual frames from movies of the same segment, colored according to normalized fluorescence (ΔF/F_0_), showing localization of stimulus-induced calcium transients **d)** Visually evoked fluorescence transients from filopodia (F1, F2) and adjacent dendritic shafts (D1, D2). Transients were evoked by alternating full-field ON and OFF illumination (bottom). After baseline probing, the tectum was infused with the NMDAR antagonist APV. Vertical lines denote the movie frames in (c). Filopodial transients are NMDAR-dependent and are greatly attenuated in the adjacent dendritic shaft. **e)** (top) Overlaid ΔF/F_0_ traces from the base (cyan) and tip (red) of the same filopodium. (bottom) Diffusion-compensated activity (navy) removes signals entering the tip via the base. **f)** Fitting of mixing coefficients for diffusion compensation. Simultaneously measured ΔF/F0 signals at the base and tip of a filopodium. Imaging sites shown at right. A mixing coefficient (red) was fit to the lower edge of this distribution for each filopodium using robust regression, to compensate for diffusing signals. Scalebar: 2μm. **g)** Mean noise correlations in all neurons measured for spontaneous (red tones) and evoked (blue tones) activity, between pairs of filopodium tips, for calcium transients (black) and diffusion-compensated signals (blue), binned by intracellular distance. Spontaneous activity is more highly correlated than evoked activity. **: p<0.01, ANCOVA followed by Tukey LSD. N=13 neurons, 267404 pairs. Error bars indicate ± SEM.

**Fig. S5.**
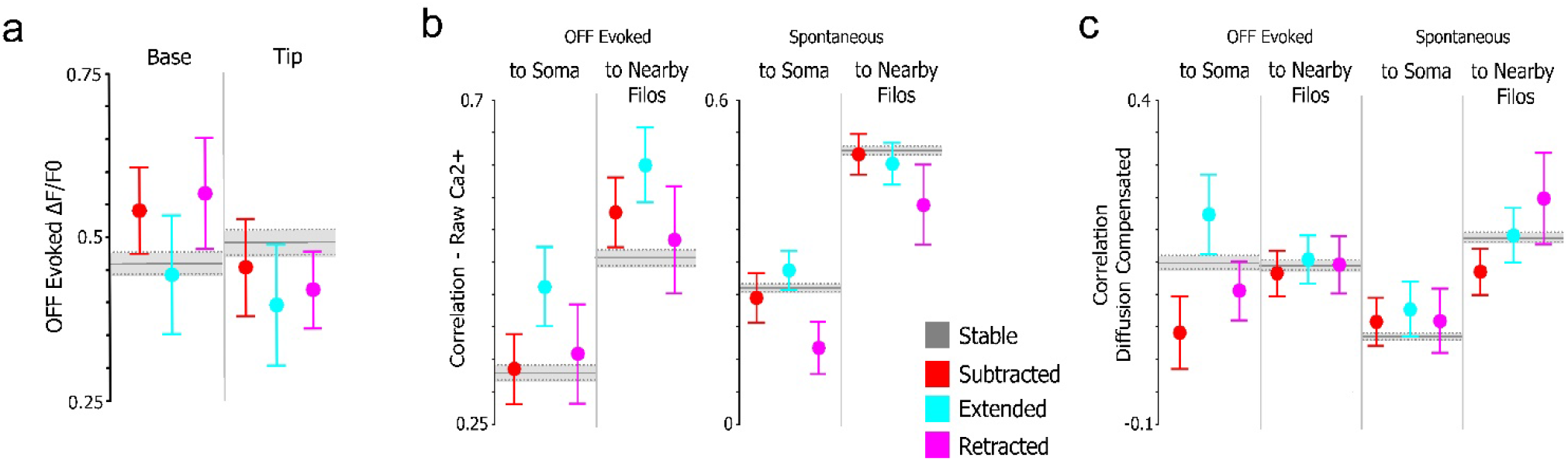
**a)** Mean pretraining OFF-evoked transient amplitudes at bases and tips of filopodia, grouped according to their motility during training. Stable filopodia values are represented by grey horizontal line. **b-c)** Correlations in activity are not strongly associated with training-induced motility in tectal neuron filopodia. Motile or retracted filopodia did not differ from stable filopodia in correlations to neighbors (<10um), or to the soma, in either evoked or spontaneous activity. N = 13 neurons, 1152 stable filopodia, 41 subtractions, 31 extensions, 25 retractions. (p > 0.05 for all, Tukey HSD compared to stable). Error bars indicate ± SEM in all subfigures.

## Notes

### Competing Interest Statement

The authors have declared no competing interest.

## References

1. T. Branco, M. Häusser, Synaptic integration gradients in single cortical pyramidal cell dendrites. Neuron. 69, 885–892 (2011).

2. T. Branco, M. Häusser, The single dendritic branch as a fundamental functional unit in the nervous system. Curr. Opin. Neurobiol. 20, 494–502 (2010).

3. T. J. McBride, A. Rodriguez-Contreras, A. Trinh, R. Bailey, W. M. Debello, Learning drives differential clustering of axodendritic contacts in the barn owl auditory system. J. Neurosci. 28, 6960–6973 (2008).

4. M. Lavzin, S. Rapoport, A. Polsky, L. Garion, J. Schiller, Nonlinear dendritic processing determines angular tuning of barrel cortex neurons in vivo. Nature. 490, 397–401 (2012).

5. H. Jia, N. L. Rochefort, X. Chen, A. Konnerth, Dendritic organization of sensory input to cortical neurons in vivo. Nature. 464, 1307–1312 (2010).

6. Z. Varga, H. Jia, B. Sakmann, A. Konnerth, Dendritic coding of multiple sensory inputs in single cortical neurons in vivo. Proc. Natl. Acad. Sci. U.S.A. 108, 15420–15425 (2011).

7. X. Chen, U. Leischner, N. L. Rochefort, I. Nelken, A. Konnerth, Functional mapping of single spines in cortical neurons in vivo. Nature. 475, 501–505 (2011).

8. T. Chen, T. J. Wardill, Y. Sun, S. R. Pulver, S. L. Renninger, A. Baohan, E. R. Schreiter, R. A. Kerr, M. B. Orger, V. Jayaraman, L. L. Looger, K. Svoboda, D. S. Kim, Ultrasensitive fluorescent proteins for imaging neuronal activity. Nature. 499, 295–300 (2013).

9. T. Kleindienst, J. Winnubst, C. Roth-Alpermann, T. Bonhoeffer, C. Lohmann, Activity-dependent clustering of functional synaptic inputs on developing hippocampal dendrites. Neuron. 72, 1012–1024 (2011).

10. N. Takahashi, K. Kitamura, N. Matsuo, M. Mayford, M. Kano, N. Matsuki, Y. Ikegaya, Locally synchronized synaptic inputs. Science. 335, 353–356 (2012).

11. S. Druckmann, L. Feng, B. Lee, C. Yook, T. Zhao, J. C. Magee, J. Kim, Structured synaptic connectivity between hippocampal regions. Neuron. 81, 629–640 (2014).

12. M. F. Iacaruso, I. T. Gasler, S. B. Hofer, Synaptic organization of visual space in primary visual cortex. Nature. 547, 449–452 (2017).

13. B. Scholl, D. E. Wilson, D. Fitzpatrick, Local Order within Global Disorder: Synaptic Architecture of Visual Space. Neuron. 96, 1127–1138.e4 (2017).

14. D. E. Wilson, D. E. Whitney, B. Scholl, D. Fitzpatrick, Orientation selectivity and the functional clustering of synaptic inputs in primary visual cortex. Nat. Neurosci. 19, 1003–1009 (2016).

15. K.-S. Lee, K. Vandemark, D. Mezey, N. Shultz, D. Fitzpatrick, Functional Synaptic Architecture of Callosal Inputs in Mouse Primary Visual Cortex. Neuron. 101, 421–428.e5 (2019).

16. M. De Roo, P. Klauser, D. Muller, LTP Promotes a Selective Long-Term Stabilization and Clustering of Dendritic Spines. PLoS Biol. 6 (2008), doi:10.1371/journal.pbio.0060219.

17. A. C. Frank, S. Huang, M. Zhou, A. Gdalyahu, G. Kastellakis, T. K. Silva, E. Lu, X. Wen, P. Poirazi, J. T. Trachtenberg, A. J. Silva, Hotspots of dendritic spine turnover facilitate clustered spine addition and learning and memory. Nature Communications. 9, 422 (2018).

18. N. Ju, Y. Li, F. Liu, H. Jiang, S. L. Macknik, S. Martinez-Conde, S. Tang, Spatiotemporal functional organization of excitatory synaptic inputs onto macaque V1 neurons. Nat Commun. 11 (2020), doi:10.1038/s41467-020-14501-y.

19. K. D. R. Sakaki, P. Coleman, T. D. Toth, C. Guerrier, K. Haas, in 2018 40th Annual International Conference of the IEEE Engineering in Medicine and Biology Society (EMBC) (2018), pp. 1–7.

20. K. D. R. Sakaki, K. Podgorski, T. A. Dellazizzo Toth, P. Coleman, K. Haas, Comprehensive Imaging of Sensory-Evoked Activity of Entire Neurons Within the Awake Developing Brain Using Ultrafast AOD-Based Random-Access Two-Photon Microscopy. Front Neural Circuits. 14, 33 (2020).

21. G. Duemani Reddy, K. Kelleher, R. Fink, P. Saggau, Three-dimensional random access multiphoton microscopy for functional imaging of neuronal activity. Nature Neuroscience. 11, 713–720 (2008).

22. D. Dunfield, K. Haas, Metaplasticity governs natural experience-driven plasticity of nascent embryonic brain circuits. Neuron. 64, 240–250 (2009).

23. J. H. Bollmann, F. Engert, Subcellular Topography of Visually Driven Dendritic Activity in the Vertebrate Visual System. Neuron. 61, 895–905 (2009).

24. K. Podgorski, D. Dunfield, K. Haas, Functional Clustering Drives Encoding Improvement in a Developing Brain Network during Awake Visual Learning. PLoS Biology. 10, e1001236 (2012).

25. W. C. Sin, K. Haas, E. S. Ruthazer, H. T. Cline, Dendrite growth increased by visual activity requires NMDA receptor and Rho GTPases. Nature. 419, 475–480 (2002).

26. M. Munz, D. Gobert, A. Schohl, J. Poquérusse, K. Podgorski, P. Spratt, E. S. Ruthazer, Rapid Hebbian axonal remodeling mediated by visual stimulation. Science. 344, 904–909 (2014).

27. Y. Mu, M.-M. Poo, Spike timing-dependent LTP/LTD mediates visual experiencedependent plasticity in a developing retinotectal system. Neuron. 50, 115–125 (2006).

28. J. Li, A. Erisir, H. Cline, In vivo time-lapse imaging and serial section electron microscopy reveal developmental synaptic rearrangements. Neuron. 69, 273–286 (2011).

29. S. Hossain, D. Sesath Hewapathirane, K. Haas, Dynamic morphometrics reveals contributions of dendritic growth cones and filopodia to dendritogenesis in the intact and awake embryonic brain. Developmental Neurobiology. 72, 615–627 (2012).

30. C. M. Niell, M. P. Meyer, S. J. Smith, In vivo imaging of synapse formation on a growing dendritic arbor. Nature Neuroscience (2004), doi:10.1038/nn1191.

31. K. Kitamura, B. Judkewitz, M. Kano, W. Denk, M. Häusser, Targeted patch-clamp recordings and single-cell electroporation of unlabeled neurons in vivo. Nat. Methods. 5, 61–67 (2008).

32. K. Haas, W. C. Sin, A. Javaherian, Z. Li, H. T. Cline, Single-cell electroporation for gene transfer in vivo. Neuron. 29, 583–591 (2001).

33. G. Y. Lazar, The development of the optic tectum in Xenopus laevis: a Golgi study. Journal of anatomy. 116, 347 (1973).

34. D. Dunfield, K. Haas, In vivo single-cell excitability probing of neuronal ensembles in the intact and awake developing Xenopus brain. Nat Protoc. 5, 841–848 (2010).

35. J. N. Bourne, K. M. Harris, Coordination of size and number of excitatory and inhibitory synapses results in a balanced structural plasticity along mature hippocampal CA1 dendrites during LTP. Hippocampus. 21, 354–373 (2011).

36. K. J. Lee, I. S. Park, H. Kim, W. T. Greenough, D. T. S. Pak, I. J. Rhyu, Motor skill training induces coordinated strengthening and weakening between neighboring synapses. J. Neurosci. 33, 9794–9799 (2013).

37. D. Smetters, A. Majewska, R. Yuste, Detecting action potentials in neuronal populations with calcium imaging. Methods. 18, 215–221 (1999).

38. B. N. Armbruster, X. Li, M. H. Pausch, S. Herlitze, B. L. Roth, Evolving the lock to fit the key to create a family of G protein-coupled receptors potently activated by an inert ligand. Proc. Natl. Acad. Sci. U.S.A. 104, 5163–5168 (2007).

39. P. Poirazi, B. W. Mel, Impact of active dendrites and structural plasticity on the memory capacity of neural tissue. Neuron. 29, 779–796 (2001).

40. R. H. Roth, R. H. Cudmore, H. L. Tan, I. Hong, Y. Zhang, R. L. Huganir, Cortical Synaptic AMPA Receptor Plasticity during Motor Learning. Neuron. 105, 895–908.e5 (2020).

41. H. Dana, Y. Sun, B. Mohar, B. K. Hulse, A. M. Kerlin, J. P. Hasseman, G. Tsegaye, A. Tsang, A. Wong, R. Patel, J. J. Macklin, Y. Chen, A. Konnerth, V. Jayaraman, L. L. Looger, E. R. Schreiter, K. Svoboda, D. S. Kim, High-performance calcium sensors for imaging activity in neuronal populations and microcompartments. Nat Methods. 16, 649–657 (2019).

42. C. Lohmann, A. Finski, T. Bonhoeffer, Local calcium transients regulate the spontaneous motility of dendritic filopodia. Nat. Neurosci. 8, 305–312 (2005).

43. C. Lohmann, K. L. Myhr, R. O. L. Wong, Transmitter-evoked local calcium release stabilizes developing dendrites. Nature. 418, 177–181 (2002).

44. M. Maletic-Savatic, R. Malinow, K. Svoboda, Rapid dendritic morphogenesis in CA1 hippocampal dendrites induced by synaptic activity. Science. 283, 1923–1927 (1999).

45. H. Makino, R. Malinow, Compartmentalized versus global synaptic plasticity on dendrites controlled by experience. Neuron. 72, 1001–1011 (2011).

46. U. V. Nägerl, N. Eberhorn, S. B. Cambridge, T. Bonhoeffer, Bidirectional activitydependent morphological plasticity in hippocampal neurons. Neuron. 44, 759–767 (2004).

47. C. Lohmann, T. Bonhoeffer, A role for local calcium signaling in rapid synaptic partner selection by dendritic filopodia. Neuron. 59, 253–260 (2008).

48. C. D. Harvey, R. Yasuda, H. Zhong, K. Svoboda, The spread of Ras activity triggered by activation of a single dendritic spine. Science. 321, 136–140 (2008).

49. U. Frey, R. G. Morris, Synaptic tagging and long-term potentiation. Nature. 385, 533–536 (1997).

50. M. E. J. Sheffield, D. A. Dombeck, Calcium transient prevalence across the dendritic arbour predicts place field properties. Nature. 517, 200–204 (2015).

51. S. X. Chen, A. Cherry, P. K. Tari, K. Podgorski, Y. K. K. Kwong, K. Haas, The Transcription Factor MEF2 Directs Developmental Visually Driven Functional and Structural Metaplasticity. Cell. 151, 41–55 (2012).

52. L. C. Andreae, J. Burrone, Spontaneous Neurotransmitter Release Shapes Dendritic Arbors via Long-Range Activation of NMDA Receptors. Cell Rep. 10, 873–882 (2015).

53. N. Spruston, Y. Schiller, G. Stuart, B. Sakmann, Activity-dependent action potential invasion and calcium influx into hippocampal CA1 dendrites. Science. 268, 297–300 (1995).

54. M. Fu, X. Yu, J. Lu, Y. Zuo, Repetitive motor learning induces coordinated formation of clustered dendritic spines in vivo. Nature. 483, 92–95 (2012).

55. K. Haas, K. Jensen, W. C. Sin, L. Foa, H. T. Cline, Targeted electroporation in Xenopus tadpoles in vivo – from single cells to the entire brain. Differentiation. 70, 148–154 (2002).

56. K. Podgorski, K. Haas, Fast non-negative temporal deconvolution for laser scanning microscopy. Journal of Biophotonics. 6, 153–162 (2013).

57. T. Laviv, B. B. Kim, J. Chu, A. J. Lam, M. Z. Lin, R. Yasuda, Simultaneous dual-color fluorescence lifetime imaging with novel red-shifted fluorescent proteins. Nat. Methods. 13, 989–992 (2016).

58. J. H. Kim, S.-R. Lee, L.-H. Li, H.-J. Park, J.-H. Park, K. Y. Lee, M.-K. Kim, B. A. Shin, S.-Y. Choi, High cleavage efficiency of a 2A peptide derived from porcine teschovirus-1 in human cell lines, zebrafish and mice. PLoS ONE. 6, e18556 (2011).

59. P. Coupé, M. Munz, J. V. Manjón, E. S. Ruthazer, D. L. Collins, A CANDLE for a deeper in vivo insight. Med Image Anal. 16, 849–864 (2012).

60. J. D. Lechleiter, D.-T. Lin, I. Sieneart, Multi-photon laser scanning microscopy using an acoustic optical deflector. Biophys. J. 83, 2292–2299 (2002).

